# Evolution of hierarchical switching pattern in antigenic variation of *Plasmodium falciparum* under variable host immunity

**DOI:** 10.1101/2023.08.30.555470

**Authors:** Gayathri Priya Iragavarapu, HJ Varsha, Shruthi Sridhar Vembar, Bhaswar Ghosh

## Abstract

*T*he var genes family encoding the variants of the erythrocyte membrane protein of *Plasmodium falciparum* is crucial for virulence of the parasite inside host. The transcriptional output of the var genes switches from one variant to other in a mutually exclusive fashion. It is proposed that a biased hierarchical switching pattern optimizes the growth and survival of the parasite inside the host. Apart from the hierarchical switching pattern, it is also well established that the intrinsic switching rates vary widely among the var genes. The centromeric protein like Var2csa is much more stable than the genes located at the telomeric and sub-telomeric regions of the chromosomes. In this study, we explored the evolutionary advantage achieved through selecting variable switching rates. Our theoretical analysis based on a mathematical model coupled with single cell RNA-seq data suggests that the variable switching rate is beneficial when cells expressing different variants are deferentially amenable to be cleared by the immune response. In fact, the variants which are cleared by the immune systems more efficiently are more stably expressed compared to a variant attacked by the immune system much less vigorously. The cells turn off expression of the variant quickly which is not cleared very efficiently. The evolutionary simulation shows that this strategy maximizes the growth of the parasite population under the presence of immune attack by the host. In corroboration with the result, we observed that stable variant has higher binding affinity to IgM from experimental data. Our study provides an evolutionary basis of widely variable switching rates of the var genes in *Plasmodium falciparum*.

## Introduction

The persistence of *Plasmodium falciparum* inside the host, clinically the most virulent Plasmodium specie, crucially relies on the switching between the variants of antigenic protein pfEMP1. During blood-stage infection, *P. falciparum* secretes a single transmembrane domain-containing protein called PfEMP1 (*P. falciparum* Erythrocyte Membrane Protein 1) which localizes to parasite-created structures called knobs on the erythrocyte membrane and triggers cytoadherence of the infected erythrocyte to the blood endothelium to avoid splenic clearance^1^. PfEMP1 also mediates the interaction of the infected erythrocyte with uninfected ones, resulting in a phenomenon called rosetting^2^. Importantly, given its abundance on the surface of the infected erythrocyte, PfEMP1 is the main target of the host immune system and antibodies are rapidly produced against it^2^. In order to evade immune detection, the parasite has evolved a strategy where it can switch between different variants of the antigenic PfEMP1 protein rendering the previously produced antibodies ineffective^3-5^. PfEMP1 is encoded by ∼60 genes of the *var* multigene family, which are located as singlets or doublets in telomeric and subtelomeric regions of thirteen of the fourteen linear *P. falciparum* chromosomes, and as tandem arrays in central regions of chromosomes 4, 6, 7, 8 and 12^6^. The structure of *var* genes consists of two exons separated by an intron (**Figure 1C**): exon 1 encodes for the outer, surface-exposed, highly variable domain of PfEMP1 and exon 2 encodes for a relatively well-conserved inner domain. The ∼60 *var* genes display a mutually exclusive expression pattern, with each individual parasite expressing a single *var* gene^4-5^. Although at a “bulk” population level, multiple *var* genes may be expressed, within a single cell, a single *var* gene is actively transcribed and its corresponding PfEMP1 protein is expressed on the infected erythrocyte surface.

**Figure 1:**
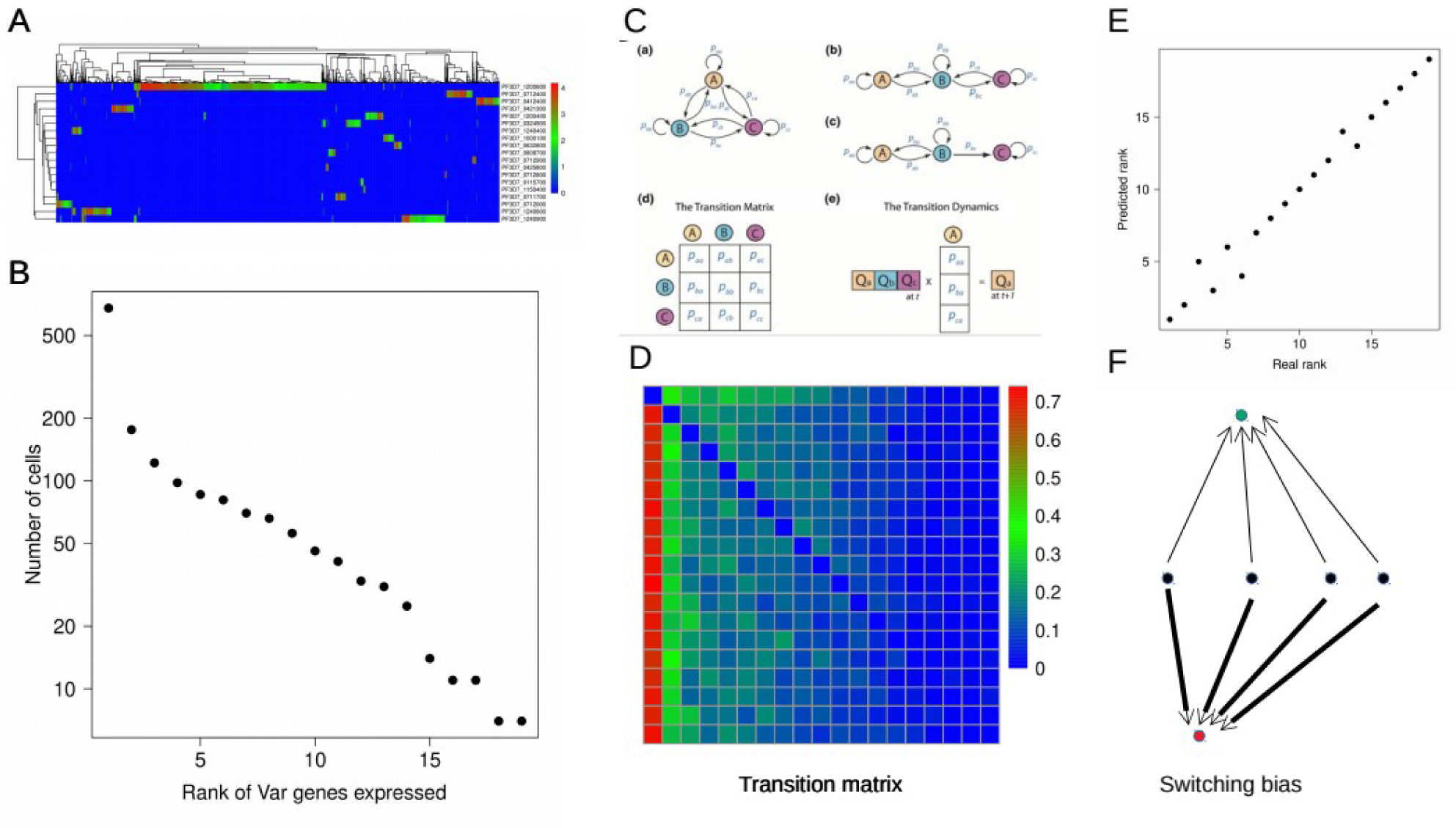
Markov Chain Transition model shows a hierarchical the transition matrix of antigenic switching. (A) The heat map displays the RPKM (Reads per Kilo base per Million reads) values of expression of the 19 expressed var genes over a population of 1515 cells. Each column represents one cell and each row represents one Var gene. The names of the Var genes are indicated. (B) The number cells expressing a particular var genes are ranked from high values to low values. (C) A schematic diagram displays the Markov Chain Transition model for 3 Var genes utilized to calculate the transition matrix. (D) The transition matrix of the probability of transition from one var gene to another is shown. The column values corresponds to the transition probability towards the particular var genes where as row values represent transition probability out of the particular var gene. The matrix columns are ordered according to the order of the hierarchy as in Figure 1B. (E) The real and the predicted rank from the model are plotted as scatter plot. (F) A schematic diagram shows the basic structure of the transition matrix in the form of a network. Nodes here represent Var genes and links correspond to the bias in transition. The arrows describes biased transition of other var genes towards the first Var gene in the hierarchy (red dot) and the second Var gene (green dot) in the hierarchy. The width of the line codes for the value of the transition bias.

Mutually exclusively *var* expression is the best studied gene regulatory mechanism in *Plasmodium* parasites. Epigenetic regulation by histone post-translational modifications (PTMs), non-coding RNAs, nuclear architecture including 3D chromatin conformation, and interaction between the promoter and intronic regions of *var* genes, are some of the key processes that ensure that a single variant gene is expressed in any given parasite (reviewed in^3,7-10^). However, the factors that drive the switch in expression pattern from one variant to the other are not fully understood. Experimental data using laboratory-adapted parasites suggested that dynamic changes in the local chromosome conformation may underlie *var* switch patterns and that different *var* genes may have intrinsically different switching rates^11-21^. Central *var* genes, in particular, appear to be expressed in a relatively stable manner, with low switching rates, while subtelomeric *var* genes appear to have higher “on” and “off” rates. Other studies combining mathematical modelling with experimentation inferred *var* gene-specific switching rates^19^, and derived a complex but coordinated pattern of switching^11,18,20^. In general, all of these studies point to a hard-wired hierarchy of *var* expression. To date, a majority of the *var* expression studies have relied on bulk transcriptomics or quantitative Reverse Transcriptase-PCR (qRT-PCR) approaches, which severely obstructs the observation of mutually exclusive expression patterns at the single cell level. With the advent of single-cell analysis methods, malaria researchers attempted different techniques such as Fluidigm C1 and Drop-seq to identify transcriptional heterogeneity within parasite populations^22-26^. These sc-RNA seq data sets can be subsequently utilized to generate the switching pattern of the Var genes in conjunction with a mathematical model previously constructed^18^.

The mathematical modeling studies suggested that the hierarchical pattern entails higher growth of the parasite inside the host in comparison to random switching^18^. However, in the sc-RNA seq data a particular hierarchy is observed where Var2csa is at the top^23^. Any other hierarchy would by symmetry exhibit the same phenomena. The particular observed pattern must have evolved due to some diversity in the immune response of the host in targeting different variants. Additionally, how the variable intrinsic switching rates among the variants would shape the evolutionary landscape is not fully explored yet. Evolutionary simulation^18^ illustrated the benefit of the hierarchical switching pattern among the variants where switching biases effectively expands the infection interval leading to higher growth rate of the population as a whole compared to the unbiased random switching. But, the intrinsic switching rates as well as the immune responses for different variants are assumed to be same for all the variants.

Here, we construct a mathematical model by incorporating response of the immune system and subsequent clearing of the variants following similar method^18^. The biased switching pattern in the absence of immune system is initially calibrated using the single cell RNA-sequencing data sets providing in vitro expression of all the variants over a population of the *Plasmodium falciparum* parasites^23^ in conjunction with recently measured switching pattern datasets^27^. Finally, the calibrated intrinsic switching pattern is utilized to construct a genetic algorithm based evolutionary simulation^18^ in presence of immune response where are varied through generations. We found that the growth rate is maximized when the killing rates of the variants by the immune response correlate negatively with the corresponding switching rates of the variants implying that the parasites expressing variants which are more efficiently being killed by the immune response tend to posses lower switching rates compared to the variants that are not being cleared away by the immune response. In fact, the genes having stable expression also tend to correlate with respect to experimental measured binding affinity of IgM^28^ to the corresponding protein, e.g, Var2csa. The genes with low switching are found to be overwhelmingly located in the centromeric regions corroborating our hypothesis that variants with low switching rates tend to have higher efficiency to be recognized and cleared by the host immune response. Our study provides a general evolutionary framework to investigate the role of antigenic variation in the spreading of parasitic infection.

## Results

### A biased switching pattern reproduces the expression pattern of the variants

First, in order to unravel the switching pattern between the different variants, we extracted the expression level of the pfEMP1 mRNAs at the single cell level from the malaria cell atlas data set ^23^. The data set in fact displays a mutually exclusive expression pattern in vitro over the population of cells (**Figure 1A**). Only 19 of the 62 variants are expressed in at least one cell in the population. The number cells expressing each variant are counted and the number shows an hierarchical expression levels (**Figure 1B**). The PF_1200600 which corresponds to the var2csa gene resides at the top of the hierarchy.

We constructed a mathematical model where the variants (i) can switch to any other variant (j) with intrinsic switching rate *w* _*ji*_ and each variant can grow with some constant rate. A schematic diagram is shown to illustrate the Markov transition model (**Figure 1C**). The network of switching pattern *β*_*ij*_ is calibrated by fitting the expression pattern of the variants from single cell RNA-seq data set (details in Materials and methods). The fitted transition matrix (**Figure 1D**) reproduces the expression level hierarchy as observed in the sc_RNA seq data for the 19 variants (**Figure 1E**). Since the experiment is performed in vitro, the *β*_*ij*_ network provides the intrinsic switching pattern in absence of immune response. All the variants are biased towards the variant which is most abundantly expressed PF_1200600) in the population and the biases toward the second one in hierarchy of expression subsequently follow the first one (**Figure 1F**). It is observed that for the dominant variant, the switching is quite unbiased to all the other variants, *i*.*e*., the switching events are almost equally probable to the other variants (**Figure 1D**). In order to check, if the hirrachy obsred in this particular data set shows resembence to other experimental data, we repeated the analysis with the Poran *et. al*. Data^25^ and found that the hierarchy matches significantly between the two data sets with a correlation of 0.66 (p-value=0.01; **Figure S1**). The hierarchical switching pattern with Var2csa at the top in fact is shown to coordinate the switching among different variants^52^.

### The particular hierarchy displays higher growth in presence of a correlated variation of the immune responses

We next incorporated both the adaptive and innate immunity in the model. The antibody is produced when a particular variant number exceeds a threshold (**Figure 2A**). The dynamics of the parasite population growth is simulated with the transition matrix obtained by calibration as in Figure 1D and keeping the immune response fixed against all the variants.. As a control, in order to reproduce the previous result, we also performed simulations with a set of unbiased transition matrix. The random switching events are found be less efficient for the parasite population since the parasite can survive longer for hierarchical switching due to longer time interval between the switching events compared to the random switching cases (Figure 2B). However, the growth rate of the parasite population does not depend on any particular hierarchy. Even if we change the hierarchy, the growth rate remains the same (the error bar in the growth rate curves correspond to different hierarchies).

**Figure 2:**
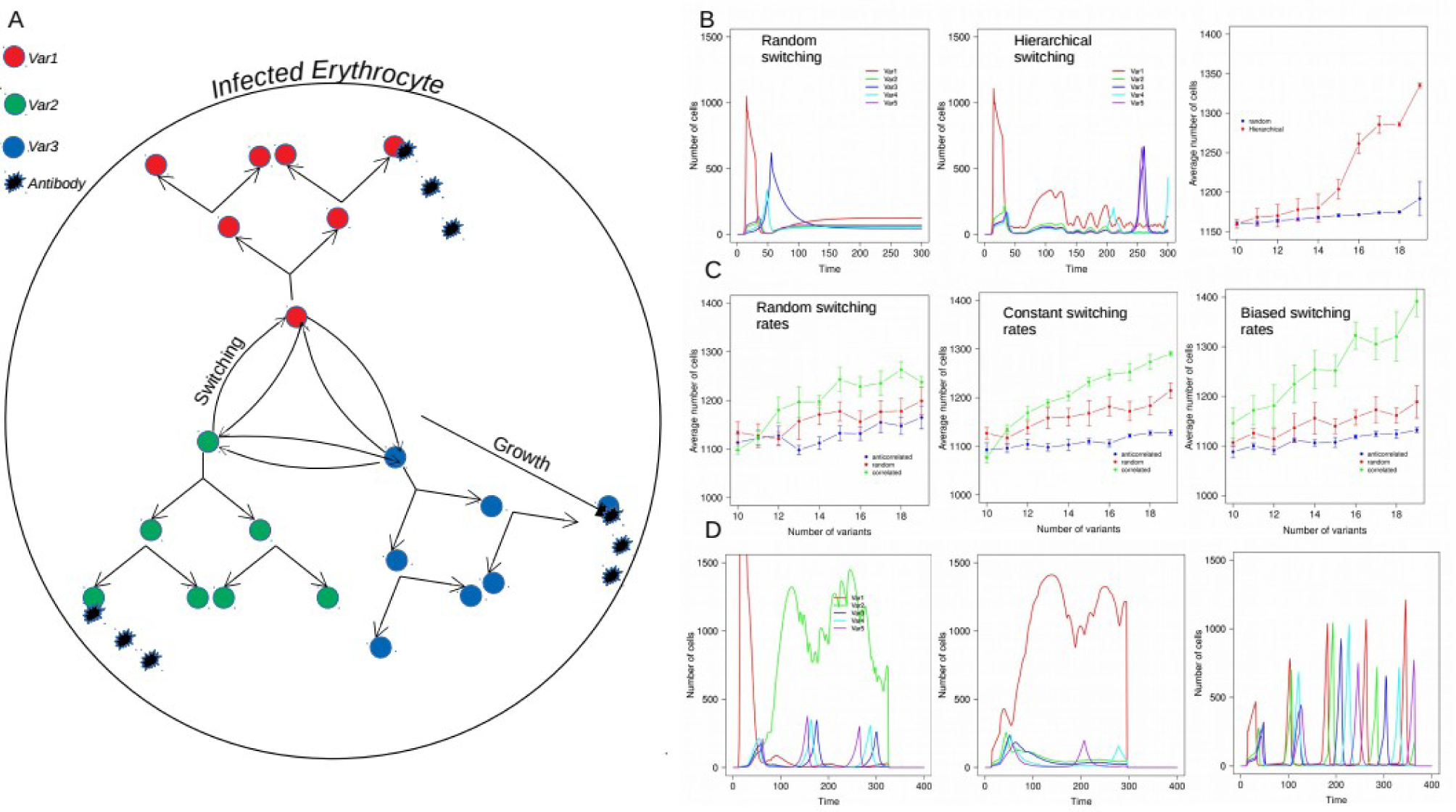
The particular hierarchical biased switching pattern is advantageous for the parasite in presence correlated bias in the immune responses. (A) The schematic diagram display the mathematical model for the host parasite interaction. The three var genes indicated by red, green and blue circles grows inside the infected erythrocyte (IE). They can switch from expressing one variant to other and are killed by the specific antibodies produced my the adaptive immunity of the host in addition to a nonspecific immune responses. The killing rate generally varies among the cells producing different Var proteins. (B) The dynamics of the number cells producing each var genes as a function of time is shown for five different types of Var genes as indicated for random switching patter (left panel), hierarchical switching pattern as calculated in Figure 1 (middle panel) and corresponding total number of cells producing all the variants for random and hierarchical switching as indicated (right panel) by varying number of existing variants. The simulation is performed with 10 different hierarchies. The graph shows the average and the error bars describe standard deviation around the average for the 100 different runs. (C) The average dynamics of the total number cells as a function of the number of existing variants by selecting the switching off rates (*s*_*i*_) random (left panel), constant (middle panel) and biased (left panel) for three different cases of correlated, anti correlated and random immune response values (*α*_*VSI*_) for adaptive immunity as indicated I the figure legend. The error bars corresponds to the standard deviation for 100 different sets of *α*_*VSI*_ values. (D) The corresponding dynamics of five variants for the biased switching rate case (Figure 2C right panel) for random (left panel), anti correlated (middle panel) and correlated (left panel) values of the adaptive immune attacks (*α*_*VSI*_). The five different variants are indicated in the figure legends.

In order to explore the effect of the particular hierarchy, we performed the simulation now by variable immune responses for different variant expressing parasites. Two possible scenarios are obvious. In one case the killing rates by the adaptive immunity is considered to be correlated with the hierarchy, meaning the dominant variant would be cleared with higher rate and in another case the killing rates are anti-correlated. As a control scenario, the killing rates are varied randomly without any preassigned correlation with the expression levels. We also select three further scenarios in the switching rates between the variants to investigate the results in presence of variable switching rates while the transition matrix remains the same. The simulations were performed by changing the number of variants from 10 to 19 to explore effect of variant number. The growth rates for the case of correlated immune responses are observed to higher irrespective of the variant number or the variation in switching rates (**Figure 2B**). In fact, the growth rate tends to increases systematically with the number of variants due to diversity in the repertoire leading to longer survival of the variants. When, the switching rates are further biased reinforcing the hierarchy, the growth rate goes to even higher level for the positively correlated case as observed in the Figure 2B, comparing the unbiased and biased switching rate scenarios. The dynamics of the different variants reveal that when the immune responses vary with the hierarchy in a positive fashion the dominant variant is cleared by the adaptive immunity faster allowing other variants also grow which leads to a longer cycle of the parasite survival. In case of anti correlated immunity, the dominant variant number grows to much higher level draining most of the finite resources required for the other variants to grow. This is also true if the immune responses are random (**Figure 2C**). Thus, one can conclude that a positively correlated hierarchy with the immune response is always preferable from the point of view of the parasite as the parasite population can sustain it’s growth for a longer time inside the host by managing the finite amount of resources in a more efficient way. Therefore, the particular hierarchy in switching pattern may have evolved in order to cater to the variable immune responses against different variants. The dominant variant var2csa would be much more efficiently cleared by the immune system compared to the less dominant one’s providing scope for all the variants to grow effectively. It is quite intuitive that the variants which is removed very efficiently do not required to produced all the time, since it will survive even if the corresponding gene is turned off, On the other hand, the variant which is immediately degraded by the immunity is required to stably expressed. As a result, this strategy elicits higher fitness to the population.

### The positively correlated hierarchy is evolutionary stable strategy

To prove whether this strategy is evolutionary stable, we next set out to perform a set of evolutionary simulations based on genetic algorithms and demonstrate that starting from a random immune response to all the variants the particular hierarchical immune response would evolve if the parasites already have the obtained hierarchically biased transition matrix. In the simulation, we start with an initial population with random immune responses and the population evolves following a genetic algorithm with parasite growth rate as the fitness parameter (**Figure 3A**, methods for details). The simulations are conducted considering three different scenarios in line with the results in Figure 2B. One scenario, where the transition matrix is hierarchical but the intrinsic switching rates are random (as in left panel of Figure 2B). In the evolutionary simulation, the population initially start at the red curve and would try to evolve towards the green curve. The simulation is performed only with 19 variants. The second scenario corresponds to the case of biased intrinsic switching rates exemplified by the right most panel of Figure 2B. As a control, the simulation is also performed for random transition probability matrix. The evolutionary dynamics clearly shows that the initial population with randomly selected immune responses slowly increase it’s fitness for both biased and unbiased intrinsic switching rates but the population with random transition matrix does not have any tendency to improve the fitness (**Figure 3B**). We also observed that for both the cases the a positive correlation between the hierarchy in switching and hierarchy in immune responses emerges (**Figure 3C**). The number of population with positive correlation also tend to increase substantially (**Figure 3D**). As expected, the population with biased intrinsic switching rate even show better performance with respect to fitness increase as well as positive correlation, although random intrinsic switching also seems to work fine albeit little slowly (**Figure 3B-D**, comparing green and red curves). These results demonstrate that the strategy proposed through the previous simulation study is indeed a evolutionary optimized stable strategy.

**Figure 3:**
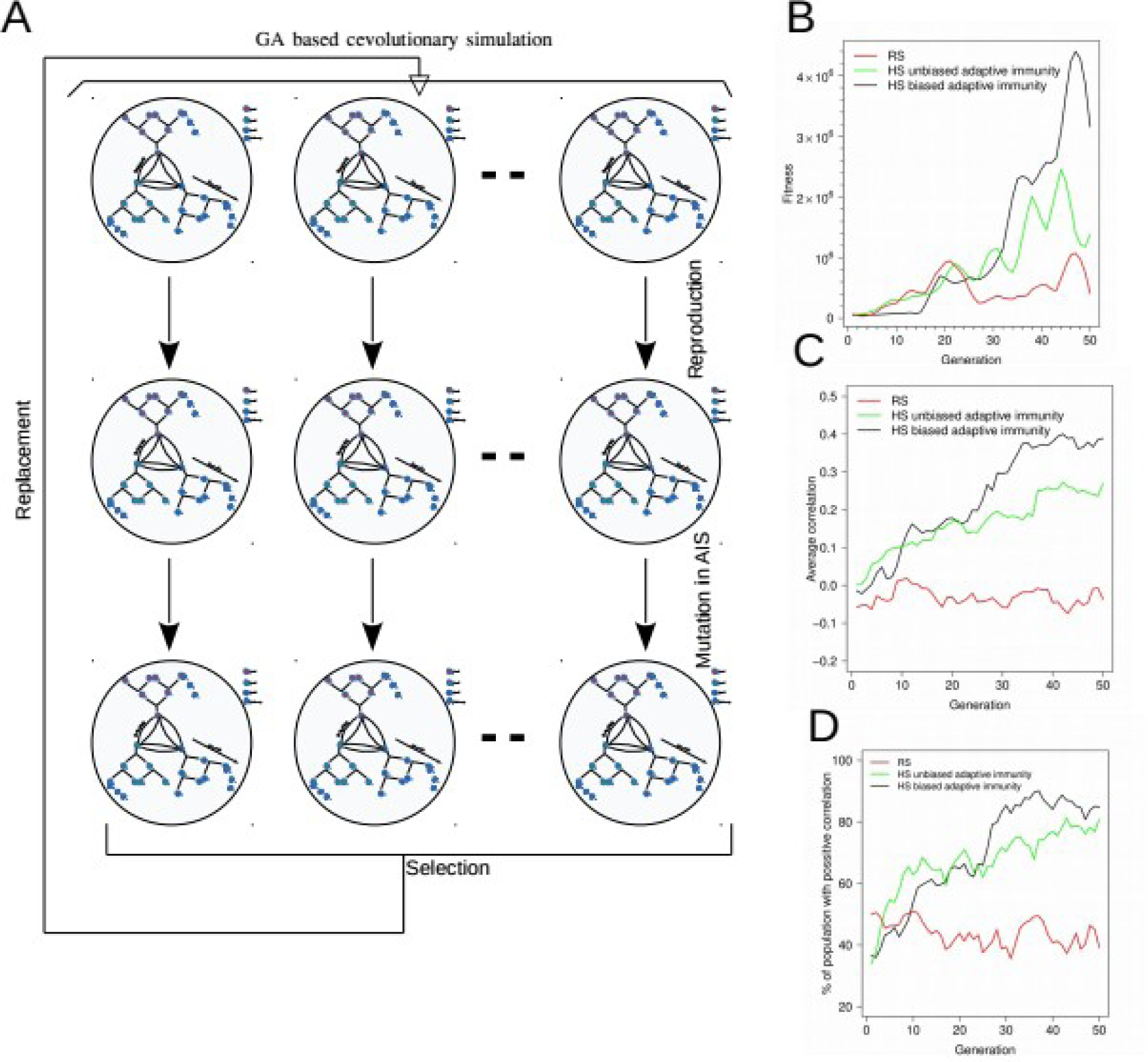
Evolutionary simulation demonstrate that the correlated adaptive immunity is an evolutionary adaptable and stable strategy. (A) A schematic diagram shows the steps in the genetic algorithm based evolutionary simulation. The initial population has random set of *σ*_*VSI*_ values for the different variants. Each generation the population is replicated and new population is selected by replacement according to the total number of parasites over time step 500. In each generation one of the *σ*_*VSI*_ is changed randomly and the process is repeated. The simulation is conducted for 100 different population of parasites over 50 generations. (B) The average fitness over 10 different runs of the parasite population as a function of generation for three different scenarios of Random transition matrix (RS), hierarchical transition matrix with unbiased switching off rates (*S*_*i*_) and hierarchical transition matrix with biased switching off rates (*S*_*i*_) as indicated in the figure legends. (C) The average correlation of the parasite population as a function of generation for three different scenarios of Random transition matrix (RS), hierarchical transition matrix with unbiased switching off rates (*S*_*i*_) and hierarchical transition matrix with biased switching off rates (*S*_*i*_) as indicated in the figure legends. (D) The average number of population with positive correlation of the parasite population as a function of generation for three different scenarios of Random transition matrix (RS), hierarchical transition matrix with unbiased switching off rates (*S*_*i*_) and hierarchical transition matrix with biased switching off rates (*S*_*i*_) as indicated in the figure legends.

### Experimental data analysis to validate the theoretical hypothesis

The hypothesis that immune response correlates with the expression hierarchy presents some obvious consequences namely the how the in vitro hierarchy would change in presence of host immunity. If the correlation was negative, then the immune response would reinforce the hierarchy essentially keeping it intact, whereas the negative correlation would tend to neutralize the hierarchy. To validate the observation, we need to investigate the expression hierarchy in presence of the host immunity. To analyze the interactions between transcriptomes of human host cells and infecting parasites, a recent study carried out RNA-seq analysis of peripheral blood samples of malaria patients^29^. They generated an average of 30 million RNA-seq tags per sample from each of 116 patients. We specifically extracted the expression levels of the 62 pfEMP1 variants for the 116 patients, The sample which has substantial number of the variants expressed are considered for further analysis. We calculated the pairwise correlation of the variants among the selected 40 patients (**Figure 4A**). The same correlation between the growth of different variants with the in vitro expression levels of the 19 variants of the sc-RNAseq data were also calculated from the simulation in presence of host immunity by sampling the dynamics for 100 times. The distribution of the correlation for three scenarios corresponding to the correlated, anti-correlated and random immune responses were estimated and compared with experimentally observed distribution of the values obtained in Figure 4A (**Figure 4B**). It is clearly observed that the experimental distribution is closest to the correlated one case. The KL-divergence also displays closer proximity to the correlated immune response case (**Figure 4C**). This demonstrates the possibility that the immune response may have been correlated with the expression level making the hierarchy less pronounced as expected from the theoretical analysis. This is an indirect validation of the proposed strategy that the genes on the top of the hierarchy would be targeted by the adaptive immunity more efficiently.

**Figure 4:**
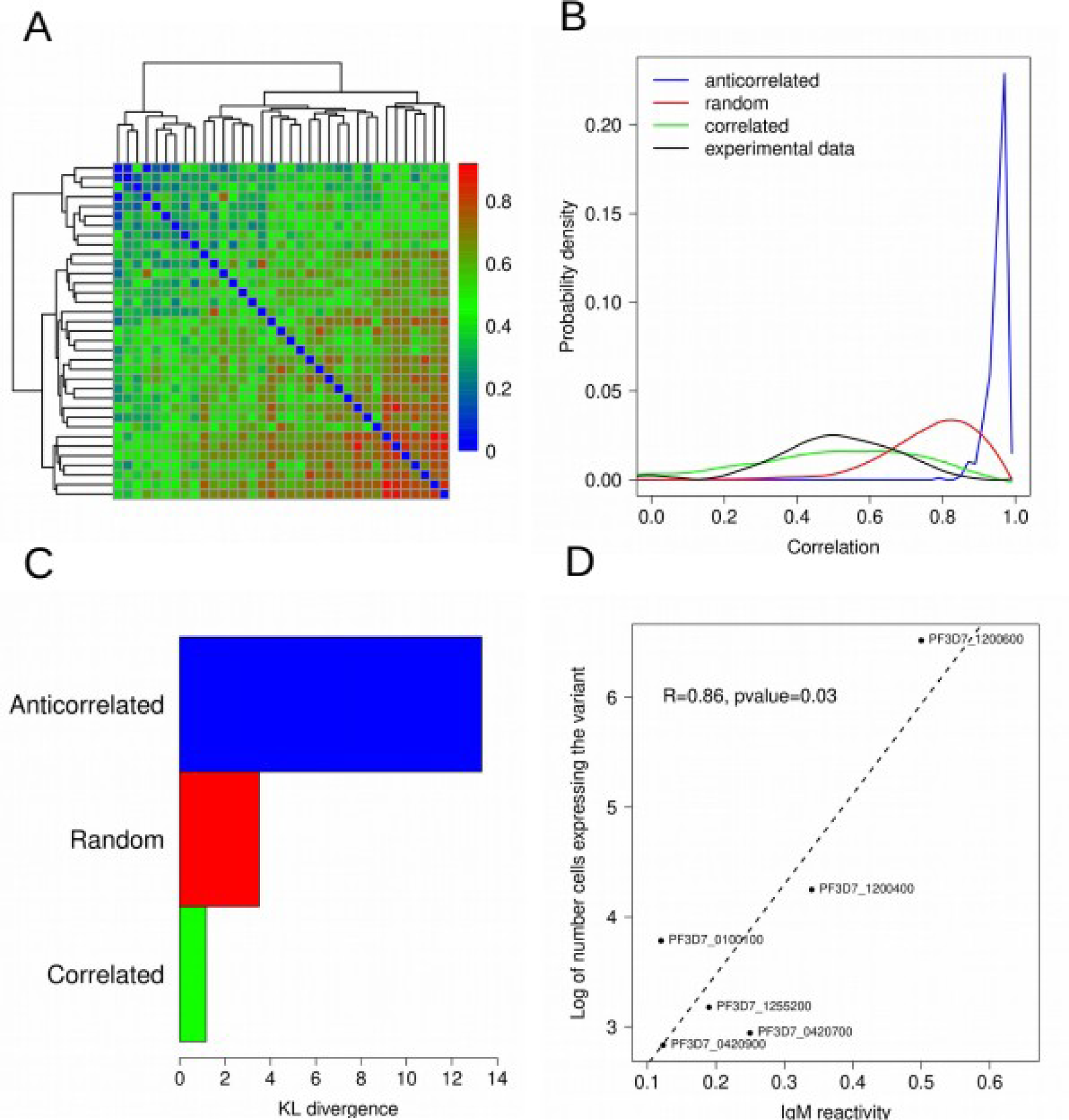
The experimental validity of the correlated adaptive immunity hypothesis. (A) The correlation matrix of var gene expression for 40 different patients. The each column and row correspond to the correlation values with other 39 patients. (B) The distribution of correlation of Var genes as quantified from the simulation for three different scenarios of random adaptive immunity values, correlated and anti correlated adaptive immunity values along with the experimentally obtained distribution from the Figure 4A data indicated in the figure legend. (C) The KL-divergence between the experimentally obtained distribution with simulated distribution for the three indicated cases. (D) The hierarchy in number cells expressing a variant vs the IgM reactivity of the corresponding variant.

In order to investigate this directly, we explored a recently published experimental data where the binding affinities of different variants to IgM were measured^28^. In corroboration of our hypothesis, the binding affinities are indeed found to be positively correlated with the hierarchy of the genes obtained from the sc-RNA seq data (**Figure 4D**).

Further, we checked the location of the genes on the chromosome and observed that the genes at the top of the hierarchy tend to be located at the centromeric region where as the genes at the bottom are mostly located at the telomeric or sub-telomeric regions. This may be responsible for the stability of the variants at the top of the hierarchy. Next epigenetic states like histone, DNA-methylation are explored through publicly available high throughput genome scale epigenetic data sets in order to delineate plausible factors involved in the stability of the expression states.

### In vitro switching rates resemble the hierarchy of the scRNA-seq data

According to our theoretical analysis (presented in Figure 1D,F), the switching bias towards a variant correlates with the hierarchy in the expression level. Thus, all the variants show maximum bias towards a variant like Var2csa (PF_120600) which is at the top of the hierarchy. In order to experimentally validate the outcome of the theoretical analysis, we explored the switching rate measurement form the recently performed experiment^27^. In this study, they utilized a transgenic strain of P. falciparum (DC-J line) where this line allows for activation of a specific var gene (PF3D7_0223500-BSD) by culturing the parasites in the presence of blasticidin-S-HCl (DC-J “ON”). As blasticidin-S-Hcl is removed, the population slowly starts to switch to other variants. The switching rate is quantified by measuring transcript levels of all the variants with time by quantitative RT-qPCR. Since, the study claims that Pf TPx-1 Is Localized to the Active var Expression Site in Ring-Stage Parasites we also look at the data where pfTpx-1 expression is knocked down by treatment of GlcN to explore if the switching rate pattern is affected. The switching rate in fact shows remarkable bias towards the PF_120600 variant in both cases after a time interval of 3 weeks (**Figure 5A,B**). We also observed a significantly strong correlation (R=-.77, p-value=0.009) between the number of cells expressing a particular variant quantified from the scRNA seq data (presented in Figure 1) and the corresponding switching rate (**Figure 5C**) even in the pfTpx-1 knockdown strain (R=−.62, p-value=0.02); **Figure 5D**). We removed the vargenes with a very low switching rates (< 0.05) from our correlation analysis. The correlation with time exhibits the over all time, the initial population takes to make the transition from the initially expressed variant to the other variants (**Figure 5E**). In addition, we also observed similar correlation pattern for other strains in which pfPtx-1 is over expressed (**Figure S2A**). var2csa levels remained low in the GFP over expression control but in that case also the pattern remains more or less the same (**Figure S2B**). The time dynamics also display similar qualitative behavior (**Figure S2C**). These result indicate that the switching pattern is remarkably robust to strain background.

**Figure 5:**
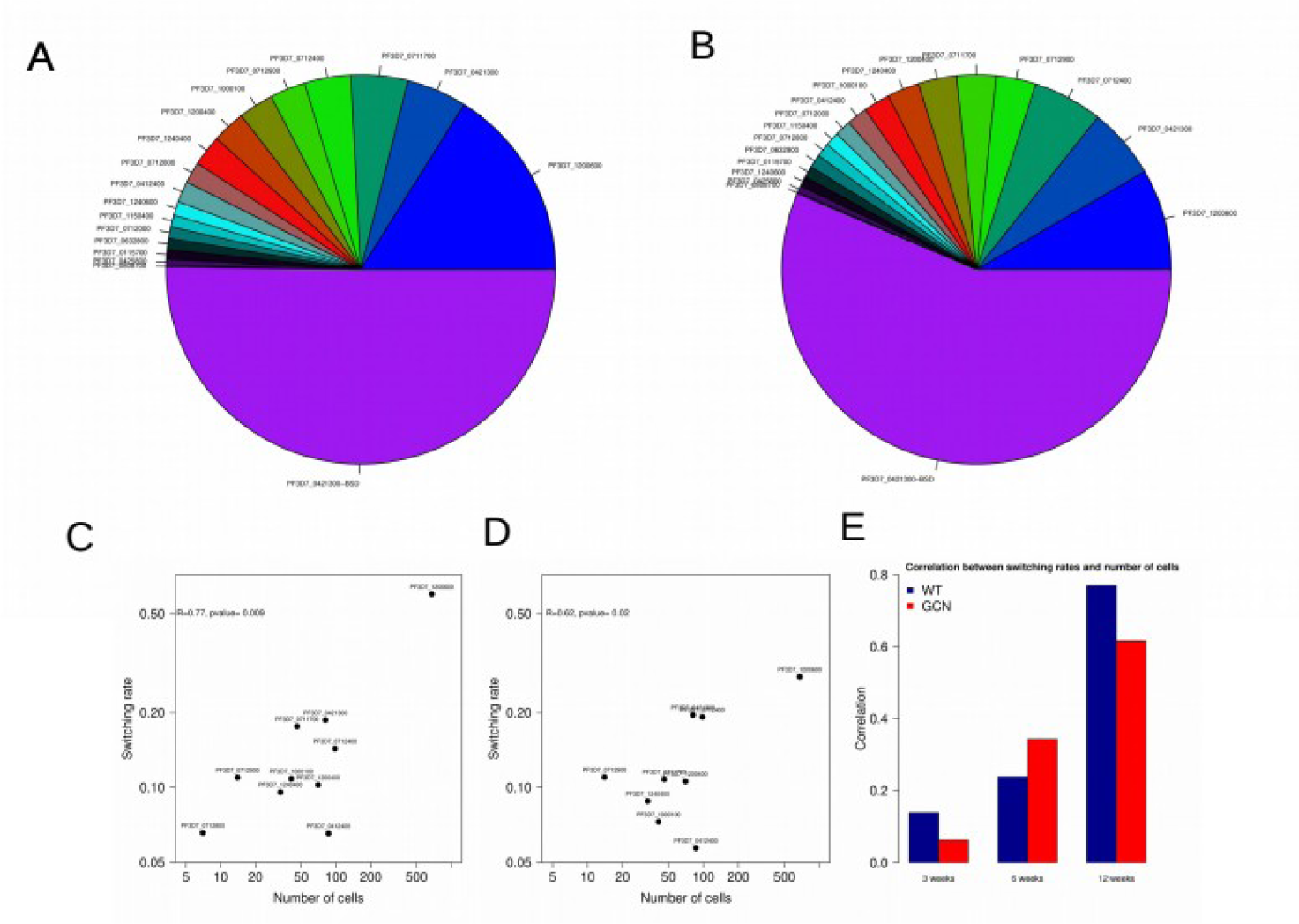
Measurement of switching rate resembles the hierarchy of expression levels of the var genes. (A) Parasites were grown in the absence or presence of 5 mM glucosamine for the duration of the experiment (±GlcN). Blasticidin was added continuously for 3 wk to induce switching to the var-bsd (+blast, red rectangle). Blasticidin was then removed from the cultures to monitor switching away from var-bsd (#blast, blue rectangle). Parasites were collected 3, 6, or 12 wk following blasticidin removal. Synchronous ring-stage parasite (20 to 22 h postsynchronization) cultures were collected for RNA, and RT-qPCR for the var gene family was performed on cDNA at the times indicated^27^. The pie chart for the off switch rates per generration of the different var genes in absence of GlcN (B) Off switch rates per generation in presence of GlcN at the time point of 3 weeks. Each pie graph represents the total of var gene transcripts with each slice of the pie representing the abundance of an individual var gene within the pool of var transcripts. (C) The scatter plot of the number of cells expressing the variants from the sc-RNA seq data and the corresponding relative mRNA expression after 3 weeks in absence of GlcN (D) The scatter plot of the number of cells expressing the variants from the sc-RNA seq data and the corresponding relative mRNA expression after 3 weeks in presence of GlcN. (E) The correlation values between the relative mRNA expression levels and the number of cells at three different times points as indicated in presence and absence of GlcN.

## Discussion

Malaria is a deadly disease caused by *Plasmodium* parasites. Approximately 226 million people are infected by malaria every year resulting in close to half a million deaths^30^. Amongst several species that cause human malaria, *Plasmodium falciparum* is the primary cause of severe infection and death, mostly of children under five years of age^31,32^. While potent drugs are available in the market for malaria treatment^33^, over the years, *Plasmodium* parasites have successfully developed resistance against many, if not all, front-line drugs^34^. This poses a serious threat to global malaria eradication efforts and the continued discovery of new drugs is necessary to tackle this debilitating disease. Eradication is further impacted by the lack of an effective malaria vaccine^35^. Despite decades of research, one of the major challenges to designing an effective blood-stage vaccine is the parasite’s remarkable ability to evade the immune response through antigenic variation^36^. Since *P. falciparum*, which adopts an obligate intracellular 48-hour life cycle within erythrocytes, is the best-studied *Plasmodium* species, we focused on antigenic variation in *P. falciparum*. Moreover, antigenic variation is a strategy for many infectious diseases^37-39^ in general which provides further motivation to understand the mechanisms and benefits of antigenic variation.

In Plasmodium species the antigenic variation of the pfEMP1 protein is a major bottleneck in designing an effective vaccine. Thus, understanding the mechanism of antigenic variation has been one of the focuses in malaria research in recent decades^3-5^. In spite of the significant efforts by the malaria research communities, the mechanism still remains elusive. All the epigenetic factors responsible for the mutually exclusive expression pattern and the intrinsic switching rates require intense exploration. Indeed lot of efforts has already been put forward in recent years. In addition to uncovering the underlying mechanisms of the antigenic switching, we also need to elicit potential evolutionary benefit incurred by the parasite population inside the host by employing different strategies related to antigenic switching. In this study, we employed an evolutionary study based on in silico genetic algorithm in order to illustrate a benefit achieved through widely variable switching rate among the variants. The selection for widely varying switching rate is specifically accelerated under a condition in which the clearance rates by the adaptive immunity also vary among the variants.

From the single cell RNA-seq data, we were able to extract the intrinsic switching pattern assuming that the variants are expressed in a mutually exclusive fashion. The assumption is only corroborated by the RNA-seq data itself. We indeed observe distinct mutually exclusive expression pattern in the single cell data. However, the mechanism of the mutually exclusive expression or switching are not incorporated in the model. In that sense, it is only a course grained phenomenological model which captures the growth of a cell depending on the variant present in the cell but the expression of the variants are not explicitly modeled. The fitness of the parasite population is simply taken to be the growth rate of the population. The simple model essentially captures the growth of the parasite inside the host in presence of adaptive and innate immunity. Details of internal machinery of the cell are ignored. It is also suggested that in addition to switching between different variants in a mutually exclusive manner, a variant also can switch between low to high expression levels^42^ which is not included in the model.

Recent experiments already show the hierarchical nature of the switching. However our analysis shows that the hierarchical switching pattern is beneficial even for random switching rates. The fundamental requirement for such switching pattern to be advantageous is also a variable clearance rate of cells expressing the different variants and we motivate the assumption based on the observation of variable binding affinities of the variants to IgM^28^. It is an indirect proof and it is not yet fully clear whether IgM is fully responsible for clearance of the variants and the role of IgM and other immunoglobulin such as IgG in the functioning of the pfmp1 protein^40,41^. More experimental studies are required to unravel the role of the specific variability of immune responses to different variants. Previous experimental observations presented the existence of highly variable immune responses among different individual patients^43,44^ but variability in immune response to different variants are not properly explored yet. Together, our analysis provides an evolutionary framework to elucidate the evolutionary optimization of switching rates in antigenic variation and hypothesize that the variable switching rate is an outcome of selection under a variable immune clearance of different variants.

In general, for parasites that produce antigenic variants within hosts, the infection continues until the host controls all variants, raises an immune response against a non-varying epitope. Antigenic variation can extend the total time before clearance^45,47^ which we also observed in our analysis. Extended infection benefits the parasite by increasing the chances for transmission to new hosts. The variant can spread again only after many resistant hosts die and are replaced by young hosts. Variants may, on the other hand, be maintained endemically in the host population. This requires a balance between the rate at which infections lead to host death or recovery and the rate at which new susceptible hosts enter the population^48^. Here we offer a evolutionary framework to study this intricate parasite-host interaction and it’s evolutionary implications. The model can further be extended incorporating a game theoretic formulation^49^.

## Materials and methods

### Analysis of single cell RNA-seq data

The count matrix of the single cell RNA-seq data from the malaria cell atlas repository^23^ were taken and the count for the 60 var genes were extracted from the count matrix table. The 60 var gene names, chromosome positions, telomere distances, promoter types information were extracted from the plasmoDB database^50^. Next, we count the number of cells in which each var gene es expressed and used for out var gene hierarchy analysis. We only choose variants which has at least 10 reads mapped onto it’s gene in one of the cells in the population.

The count matrix was taken from Poran A *et. al*. And only first three time points (30 and 36 hours) were taken since it is in general known that the pfEMP1 var genes are expressed predominantly in the ring-stage. In order to take correlation with the malaria cell atlas data set, we similarly counted the number of cells in which a particular var gene expressed. WE selected the var genes which is expressed in at least 1% pf the population in both cases.

### Mathematical model for the host-parasite interaction

We follow the model similar to the previous study^18^ (**Figure 2A**). In the model, the P.falciparum population grows inside the RBC. Each parasite cell can express only one variant and can switch from one variant (i) to another variant (j) following a probability transition matrix *T* _*ji*_. Each variant (i) posses an intrinsic off rate *s*_*i*_. Then the parasite number producing a certain variant i can be written as

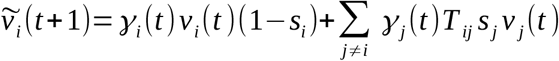

The variant gowth rate *γ*_*i*_(*t*) depends on the availability of the RBCs. Thus the growth rate at a particular time is derived from the equation

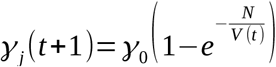

Here, *γ*_0_ is a base growth rate of the parasite population initially when the resources are plenty. N is the total number of the RBCs and 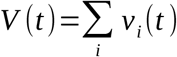 is the total number of parasites at time t.

Thus, the effective growth rate would reduce as total number of parasite increases and would tend to zero as total number is very high compared to the capacity N. This reflects the limitation in resources for the parasite. The control and removal of parasitised cells is assumed to be multi-factorial, being initially controlled by a non-specific immune response, NSI, before the variant specific immune responses (e.g. antibodies), VSI and CSI, are activated after some delay τ. The dynamics of the specific responses are given as:

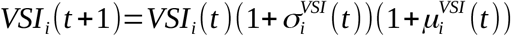

where the first terms on the r.h.s. denote immune expansion in the presence of antigen and the second terms the decay of immune cells in the absence of stimulation. The respective rates, and *σ*_*i*_ and *μ*_*i*_, are defined as

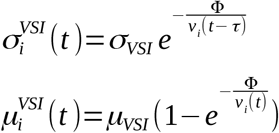

Here, *σ*_*VSI*_ and *μ*_*VSI*_ are the maximum rates of clonal expansion and decay over a 48h period; Φ is the antigen threshold levels necessary for immune stimulation, and τ is the delay of the adaptive immune response. The specific response VSI is triggered only by the presence of antigenic variant i.

With these we can now define the rate of parasite removal by the specific and cross-reactive responses, This particular form takes into consideration two important aspects: (i) the removal rate is maximized at very high levels of circulating immune cells, and (ii) the removal rate decreases as the antibody-antigen ratio declines.

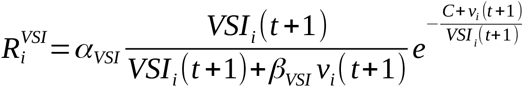

The non-adaptive immune response is thought to be immediate (within the 48h-time course of one generation) and is a simple function of parasite load, given as

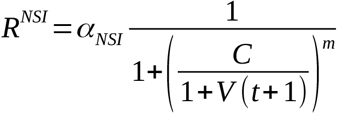

The equation for the dynamics of variant i then becomes

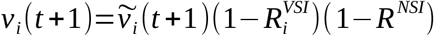

In the simulation, the initial population has the same set of randpm values the immune response represented by the parameter *σ*_*VSI*_. The values are taken to be different for different variants. For correlated case the values of *σ*_*VSI*_ are ordered from high to low following a correlation with the expression hierarchy as in Figure 1B. For anticorrelated case the values of *σ*_*VSI*_ are ordered from low to high following an anticorrelation with the expression hierarchy as in Figure 1B. For random case the values of *σ*_*VSI*_ are taken randomly.

### Metropolis algorithm for calibrating the simulation data with the scRNA-seq data

In order to calibrate the model with the scRNA-seq data, we utilized a Markov state transition model schematically described in Figure 1C. Since there are 19 var genes expressed in the data set, transition matrix as a 19 by 19 dimensional matrix. We start with a random matrix and in each time step each cell in the initial population of 100 cells can switch to one of the other 18 variants following the transition matrix. At the end of 1000 time steps, the number cells producing each variant is counted and a correlation is calculated with the experimentally observed distribution as in Figure 1B. The correlation provide us with a energy parameter (E) for the simulated annealing protocol. Then we calculate a fitness parameter exp(−E/KT) with KT=1 and use metropolis algorithm^51^ to optimize the choice of the transition matrix which fits the experimental data. We run the algorithm 500 times to find the average transition matrix.

### Genetic algorithm based evolutionary simulation

In order to evolve the population of parasites under selection for growth, a set of genetic algorithm based evolutionary simulations were performed^18^. The transition matrix obtained by fitting from the previous section is used to explore different scenarios of random immune response and biased immune response. Figure 3A shows a schematic of the evolutionary simulation protocol. The initial population has random set of *σ*_*VSI*_ values for the different variants. Each generation the population is replicated and new population is selected by replacement according to the total number of parasites over time step 500. In each generation one of the *σ*_*VSI*_ is changed randomly and the process is repeated. The simulation is conducted for 100 different population of parasites over 50 generations. Thus, the initial population would start with a very low correlation between the *σ*_*VSI*_ values and expression hierarchy and evolve towards the desired *σ*_*VSI*_ values. We repeated the evolutionary dynamics for 10 different initial samples and calculated the average dynamics for the analysis.

---

**Figure S1.**
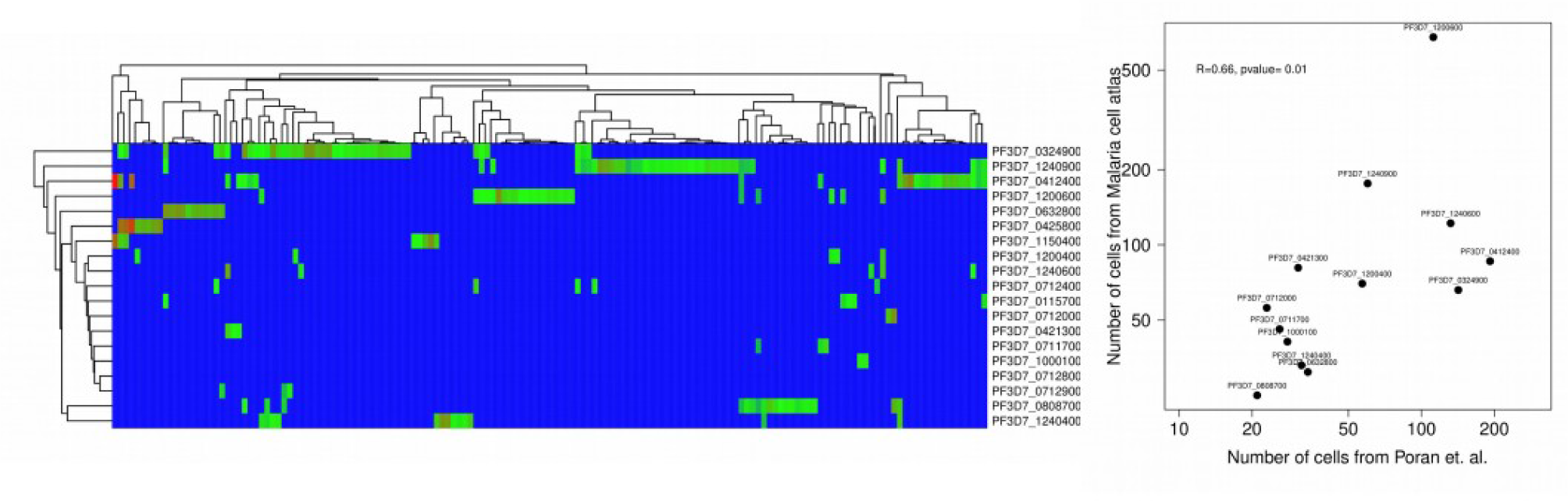

**Figure S2.**
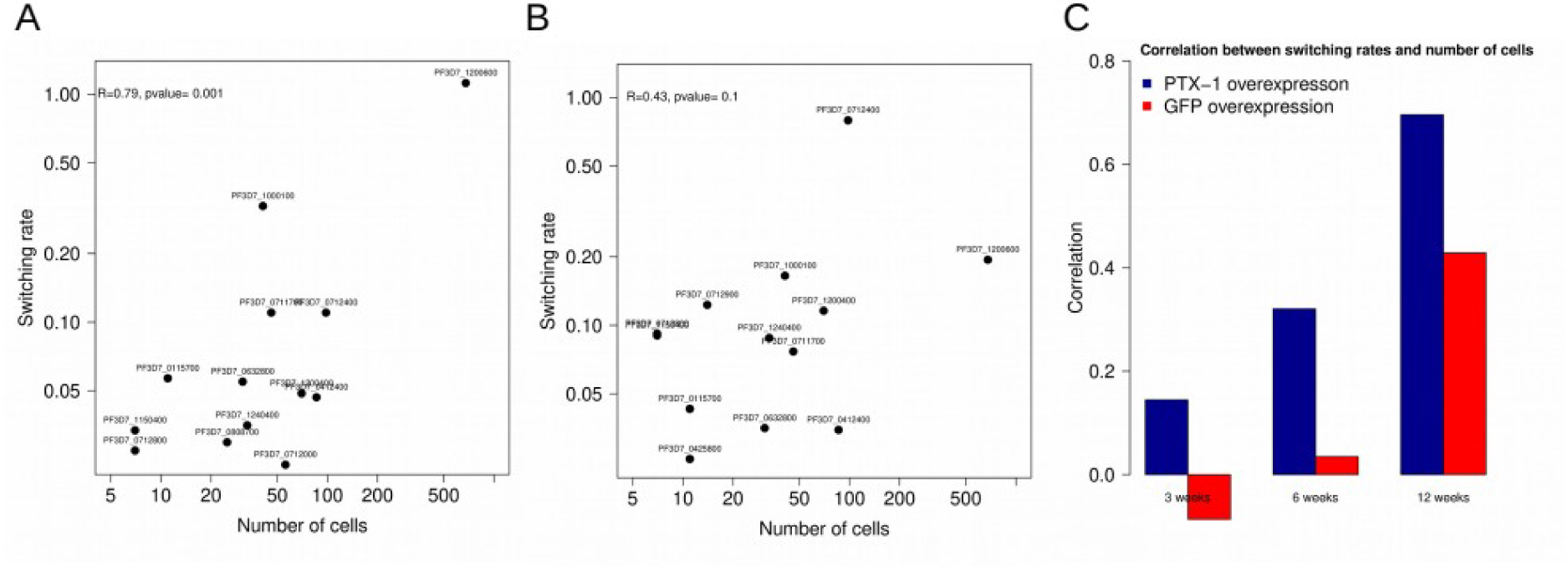

## Notes

### Competing Interest Statement

The authors have declared no competing interest.

### Summary of Updates

New authors are added and affiliations updated.

## References

1. Maier, A. G., Cooke, B. M., Cowman, A. F. & Tilley, L. Malaria parasite proteins that remodel the host erythrocyte. Nature Reviews Microbiology vol. 7 (2009).

2. Hviid, L. & Jensen, A. T. R. PfEMP1 - a parasite protein family of key importance in plasmodium falciparum malaria immunity and pathogenesis. Adv. Parasitol. 88, (2015).

3. Scherf, A., Lopez-Rubio, J. J. & Riviere, L. Antigenic variation in Plasmodium falciparum. Annual Review of Microbiology vol. 15 (2008).

4. Guizetti, J. & Scherf, A. Silence, activate, poise and switch! Mechanisms of antigenic variation in Plasmodium falciparum. Cellular Microbiology vol. 15 (2013).

5. Deitsch, K. W. & Dzikowski, R. Variant Gene Expression and Antigenic Variation by Malaria Parasites. Annu. Rev. Microbiol. 71, (2017).

6. Bahl, A. et al. PlasmoDB: The Plasmodium genome resource. A database integrating experimental and computational data. Nucleic Acids Research vol. 31 (2003).

7. Duraisingh, M. T. & Skillman, K. M. Epigenetic Variation and Regulation in Malaria Parasites. Annual Review of Microbiology vol. 72 (2018).

8. Hollin, T., Gupta, M., Lenz, T. & Le Roch, K. G. Dynamic Chromatin Structure and Epigenetics Control the Fate of Malaria Parasites. Trends in Genetics vol. 37 (2021).

9. Gupta, A. P. & Bozdech, Z. Epigenetic landscapes underlining global patterns of gene expression in the human malaria parasite, Plasmodium falciparum. International Journal for Parasitology vol. 47 (2017).

10. Deshmukh, A. S., Srivastava, S. & Dhar, S. K. Plasmodium falciparum : Epigenetic control of var gene regulation and disease. Subcell. Biochem. 61, (2013).

11. Blomqvist, K. et al. var gene transcription dynamics in Plasmodium falciparum patient isolates. Mol. Biochem. Parasitol. 170, (2010).

12. Horrocks, P., Pinches, R., Christodoulou, Z., Kyes, S. A. & Newbold, C. I. Variable var transition rates underlie antigenic variation in malaria. Proc. Natl. Acad. Sci. U. S. A. 101, (2004).

13. Fastman, Y., Noble, R., Recker, M. & Dzikowski, R. Erasing the epigenetic memory and beginning to switch-the onset of antigenic switching of var genes in plasmodium falciparum. PLoS One 7, (2012).

14. Dimonte, S. et al. Sporozoite Route of Infection Influences in Vitro var Gene Transcription of Plasmodium falciparum Parasites from Controlled Human Infections. in Journal of Infectious Diseases vol. 214 (2016).

15. Ye, R. et al. Transcription of the var genes from a freshly-obtained field isolate of Plasmodium falciparum shows more variable switching patterns than long laboratory-adapted isolates. Malar. J. 14, (2015).

16. Noble, R. et al. The antigenic switching network of Plasmodium falciparum and its implications for the immuno-epidemiology of malaria. Elife 2013, (2013).

17. Scherf, A. et al. Antigenic variation in malaria: In situ switching, relaxed and mutually exclusive transcription of var genes during intra-erythrocytic development in Plasmodium falciparum. EMBO J. 17, (1998).

18. Recker, M. et al. Antigenic variation in Plasmodium falciparum malaria involves a highly structured switching pattern. PLoS Pathog. 7, (2011).

19. Milne, K. et al. Mapping immune variation and var gene switching in naive hosts infected with Plasmodium falciparum. Elife 10, (2021).

20. Bachmann, A. et al. Highly co-ordinated var gene expression and switching in clinical Plasmodium falciparum isolates from non-immune malaria patients. Cell. Microbiol. 13, (2011).

21. Gurarie, D. et al. Mathematical modeling of malaria infection with innate and adaptive immunity in individuals and agent-based communities. PLoS One 7, (2012).

22. Ngara, M. et al. Exploring parasite heterogeneity using single-cell RNA-seq reveals a gene signature among sexual stage Plasmodium falciparum parasites. Exp. Cell Res. 371, (2018).

23. Howick, V. M. et al. The malaria cell atlas: Single parasite transcriptomes across the complete Plasmodium life cycle. Science (80-.). 365, (2019).

24. Rawat, M., Srivastava, A., Gupta, I. & Karmodiya, K. Single Cell RNA-sequencing reveals cellular heterogeneity, stage transition and antigenic variation during stress adaptation in synchronized Plasmodium falciparum. bioRxiv (2019) doi:10.1101/752543.

25. Poran, A. et al. Single-cell RNA sequencing reveals a signature of sexual commitment in malaria parasites. Nature 551, (2017).

26. Walzer, K. A., Fradin, H., Emerson, L. Y., Corcoran, D. L. & Chi, J. T. Latent transcriptional variations of individual Plasmodium falciparum uncovered by single-cell RNA-seq and fluorescence imaging. PLoS Genet. 15, (2019).

27. Heinberg A, Amit-Avraham I, Mitesser V, Simantov K, Goyal M, Nevo Y, Kandelis-Shalev S, Thompson E, Dzikowski R. A nuclear redox sensor modulates gene activation and var switching in Plasmodium falciparum. Proc Natl Acad Sci U S A. (2022).

28. Quintana, M.d.P., Ecklu-Mensah, G., Tcherniuk, S.O. et al. Comprehensive analysis of Fc-mediated IgM binding to the Plasmodium falciparum erythrocyte membrane protein 1 family in three parasite clones. Sci Rep 9, 6050 (2019).

29. Yamagishi J, Natori A, Tolba ME, Mongan AE, Sugimoto C, Katayama T, Kawashima S, Makalowski W, Maeda R, Eshita Y, Tuda J, Suzuki Y. Interactive transcriptome analysis of malaria patients and infecting Plasmodium falciparum. Genome Res. 24(9):1433–44 (2014)

30. WHO. World Malaria Report 2020. https://www.who.int/news-room/fact-sheets/detail/malaria (2020).

31. Perkins, D. J. et al. Severe malarial anemia: Innate immunity and pathogenesis. International Journal of Biological Sciences vol. 7 (2011).

32. Perlmann, P. & Troye-Blomberg, M. Malaria blood-stage infection and its control by the immune system. Folia Biologica vol. 46 (2000).

33. Tse, E. G., Korsik, M. & Todd, M. H. The past, present and future of anti-malarial medicines. Malaria Journal vol. 18 (2019).

34. Haldar, K., Bhattacharjee, S. & Safeukui, I. Drug resistance in Plasmodium. Nature Reviews Microbiology vol. 16 (2018).

35. Draper, S. J. et al. Malaria Vaccines: Recent Advances and New Horizons. Cell Host and Microbe vol. 24 (2018).

36. Servín-Blanco, R., Zamora-Alvarado, R., Gevorkian, G. & Manoutcharian, K. Antigenic variability: Obstacles on the road to vaccines against traditionally difficult targets. Human Vaccines and Immunotherapeutics vol. 12 (2016).

37. Deitsch KW, Lukehart SA, Stringer JR. Common strategies for antigenic variation by bacterial, fungal and protozoan pathogens. Nat Rev Microbiol. 2009 Jul;7(7):493–503

38. Mosa AI Antigenic Variability. Front. Immunol. 11:2057.(2020)

39. der Woude MW, Bäumler AJ. Phase and antigenic variation in bacteria. Clin Microbiol Rev. 17(3):581–611 (2004).

40. Stevenson L, Huda P, Jeppesen A, Laursen E, Rowe JA, Craig A, Streicher W, Barfod L, Hviid L. Investigating the function of Fc-specific binding of IgM to Plasmodium falciparum erythrocyte membrane protein 1 mediating erythrocyte rosetting. Cell Microbiol. 17(6):819–31 (2015).

41. Jeppesen A, Ditlev SB, Soroka V, Stevenson L, Turner L, Dzikowski R, Hviid L, Barfod L. Multiple Plasmodium falciparum Erythrocyte Membrane Protein 1 Variants per Genome Can Bind IgM via Its Fc Fragment Fcμ. Infect Immun. 83(10):3972–81 (2015).

42. Abdi AI, Warimwe GM, Muthui MK, Kivisi CA, Kiragu EW, Fegan GW, Bull PC. Global selection of Plasmodium falciparum virulence antigen expression by host antibodies. Sci Rep. 25;6:19882 (2016)

43. Wichers JS, Tonkin-Hill G, Thye T, Krumkamp R, Kreuels B, Strauss J, von Thien H, Scholz JA, Smedegaard Hansson H, Weisel Jensen R, Turner L, Lorenz FR, Schöllhorn A, Bruchhaus I, Tannich E, Fendel R, Otto TD, Lavstsen T, Gilberger TW, Duffy MF, Bachmann A. Common virulence gene expression in adult first-time infected malaria patients and severe cases. Elife 28;10:e69040 (2021).

44. Bachmann A, Bruske E, Krumkamp R, Turner L, Wichers JS, Petter M, Held J, Duffy MF, Sim BKL, Hoffman SL, Kremsner PG, Lell B, Lavstsen T, Frank M, Mordmüller B, Tannich E. Controlled human malaria infection with Plasmodium falciparum demonstrates impact of naturally acquired immunity on virulence gene expression. PLoS Pathog. 11;15(7):e1007906 (2019).

45. Deitsch KW, Moxon ER,Wellems TE. Shared themes of antigenic variation and virulence in bacterial, protozoal, and fungal infections. Microbiol Mol Biol Rev 61:281–93 (1997)

46. Fussenegger M. Different lifestyles of human pathogenic procaryotes and their strategies for phase and antigenic variation. Symbiosis 1997;22:85–153.

47. Moxon ER, Rainey PB, Nowak MA, Lenski RE. Adaptive evolution of highly mutable loci in pathogenic bacteria. Curr Biol 1994;4:24–33.

48. Steven A. Frank Immunology and Evolution of Infectious Disease Princeton University Press Princeton (2002).

49. Joel E. Cohen and Charles M. Newman Host-Parasite Relations and Random Zero-Sum Games: The Stabilizing Effect of Strategy Diversification The American Naturalist 133 533–552 (1989)

50. Bahl A, Brunk B, Crabtree J, Fraunholz MJ, Gajria B, Grant GR, Ginsburg H, Gupta D, Kissinger JC, Labo P, Li L, Mailman MD, Milgram AJ, Pearson DS, Roos DS, Schug J, Stoeckert CJ Jr, Whetzel P. PlasmoDB: the Plasmodium genome resource. A database integrating experimental and computational data. Nucleic Acids Res. 31(1):212–5 (2003)

51. Oliver Catonim Metropolis, Simulated Annealing, and Iterated Energy Transformation Algorithms: Theory and Experiments Journal of Complexity 12, 595–623 (1996)

52. Zhang X, Florini F, Visone JE, Lionardi I, Gross MR, Patel V, Deitsch KW. A coordinated transcriptional switching network mediates antigenic variation of human malaria parasites. Elife. 2022 Dec 14;11:e83840.

